# Modeling by artificial neural networks of silver carp (*Hypophthalmichthys molitrixi*) with sous vide processing on the effects of storage and processing temperatures on the microbiological status

**DOI:** 10.1101/2021.01.26.428224

**Authors:** Seyed Vali Hosseini, Milad Pero, Reza Tahergorabi, Shirin Kazemzadeh, Ricardo Santos Alemán, Jhunior Abrahan Marcia Fuentes, Ismael Montero Fernández, Lilian Sosa, Xesus Feas Sanchez

## Abstract

To evaluate and anticipate the microbial changes of silver carp (Hypophthalmichthys molitrixi) during cold storage (0, 5, 10, 15 & 21 day) at different sous vide processing temperatures (60, 65, 70, and 75 °C), changes in microbial load of Enterobacteriaceae, Lactic Acid bacteria (LAB), Pseudomonas, Psychrotrophs, and total viable count (TVC) were considered. A radial basis function neural network (RBFNN) model was established to predict the changes in the microbial content of silver carp. The critical temperature for inactivation of Enterobacteriaceae and lactic acid bacteria was 65 °C and for Pseudomonas and Psychrotrophs was 70 °C and the highest value (75 °C) was observed for the total viable count. In samples processed at 75 °C, the presence of Enterobacteriaceae, Pseudomonas and Psychrotrophs were not detectable up to 15 days of storage and lactic acid bacteria were not detectable even at the end of the storage period. The optimal ANN topology for modeling Enterobacteriaceae, Pseudomonas, and Psychrotroph contained 9 neurons in the hidden layer, but for TVC and LAB, it was 14 neurons.

## Introduction

Silver carp popularity as a source of food has been notable and this aquatic fish is also used to prevent and clear algal blooms. The silver carp has become popular worldwide because does not require supplementary feed. As a result, silver carp (*Hypophthalmichthys molitrix*) is one of the most cultured fish species in the world due to its desirable properties such as high nutritional content, fast growth, high feed efficiency, and easy cultivation [1]. Unfortunately, the food safety and spoilage of fishery products is a huge concern because of microbial cross-contamination from various sources. As a consequence, because of its neutral pH and high water content, the shelf-life of fish declines rapidly. All of these aspects enhance the need to increase the shelf life and many preservation applications have been taken into consideration including sous-vide.

Sous vide is a French word meaning “under vacuum”, and sous vide cooking is a processing method in which vacuum-packed foods in heat resistant pouches are cooked under controlled temperature and time [2]. The advantages of sous vide cooked products are that these products are not preserved by low water activity or pH and do not contain preservatives. The food safety of these types of products relies on high heat treatment and cold storage [3]. In this technique, the products are immersed in a hot water/steam oven for a longer time than conventional cooking followed by immediate cooling to ≤ 4 °C [4]. Recently, sous-vide processing was applied and combined with irradiation for mackerel fillets [5]. They concluded that the microbial count of mesophilic and psychrophilic treated by sous vide with irradiation never exceeded the standard limit. The sous-vide processing of ready-to-cook seer fish steaks was optimized by response surface methodology, and they reported the optimized process condition as 3.5% salt concentration, process temperature of 89 °C, and cooking time of 13.5 min. Processed samples at these optimized conditions were preserved up to 65 days of storage at ≤ 4 °C [6].

Modeling and optimization play an important role in the reduction of waste, product quality improvement, and profitability. As there are many modeling and optimization techniques applied in food technology [7], artificial neural networks (ANNs) have gained special attention due to their ability in modeling complex processes where there is a nonlinear relationship between the dependent and independent variables [8]. Furthermore, this modeling technique has successfully been applied in modeling and optimization of food processes [9][10][11] and microbial analysis [12][13][14][15].

As a result, the main objective of this study was to analyze the effect of sous vide processing on the microbial quality of silver carp. The steps of this research were as follows: (1) sous vide processing of silver carp at different process conditions; (2) analyzing the effect of process conditions on the microbial quality (Enterobacteriaceae, Pseudomonas, Psychrotrophs, lactic acid bacteria, and total viable count) of silver carp; (3) behavior of microbial growth in sous-vide cooked fish during cold storage (21 days); (4) modeling and simulation of microbial count by artificial neural networks.

## Material and methods

Fifteen sacrisfied silver carp sample (average weight and length of 790±20 g and 283±16 mm, respectively) were obtained from a local warm-water fish farm located in Rasht, in the north of Iran. The fish were not starved and feed commercial fish food from the Faradaneh Company, Tehran, Iran. All microbial media such as Plate count agar (PCA), Pseudomonas Agar Base, Violet red bile glucose agar, Man Rogosa Sharp Agar medium cultures were supplied by HiMedia (HiMedia Co., India).

### Sample preparation

Fish samples were cut into fillets with dimensions of 10×5×2 cm. Then, each fillet was vacuum packed by a vacuum packaging machine (Guater Control Back Co, Iran) with 95% of vacuum.

The bags used for this purpose were polyamide bag (S-Gruppen, Vinterbro, Norway) with 75 μm thickness and an oxygen transmission rate of 30 cm^3^ m^−2^ day^−1^ atm^−1^. Vacuum packed samples were cooked at 60, 65, 70, and 75 °C for 15 min in a water bath (Memmrt Co., Schwabach, Germany). After cooking, they were immediately immersed in the ice bath and then stored in the refrigerator (4 °C) for 21 days.

### Microbial analysis

Microbial analysis was carried out on both control and treated samples at 0, 3, 7, 14, and 21 days of cold storage (4°C). At each storage interval, 5 g of sample was aseptically removed with the aid of a sterile scalpel and placed in a stomacher bag. 45 mL of sterile peptone saline solution (4°C) was poured into the stomacher bag containing the sample and the mixture was homogenized using a stomacher

Plate count agar (PCA) was used for determining the total viable count by incubating at 37 °C for 48 h [16]. Pseudomonas count was carried out by using Pseudomonas Agar Base according to the procedure described by the American Public Health Association [17]. Violet red bile glucose agar was used for Enterobacteriaceae count [18]. For the determination of lactic acid bacteria, Man Rogosa Sharp Agar was applied [18]. Psychrotrophs was determined on PCA followed by incubation at 4 °C for 10 days.

### Artificial neural networks (ANNs) modeling

The experimental data were used for developing ANNs where the independent variables were temperature (°C) and storage period (day) and the dependent variable was a microbial count (Log CFU g^−1^). For developing ANNs, 70, 15, and 15%of experimental data were randomly selected for training, cross-validation, and testing, respectively. Multilayer perceptron (MLP) and ANNs were used for modeling the microbial count (Fig. 1) In the procedure of finding the best ANN structure, a different number of neurons (1 to 15) in the hidden layer was analyzed.

**Fig. 1.**
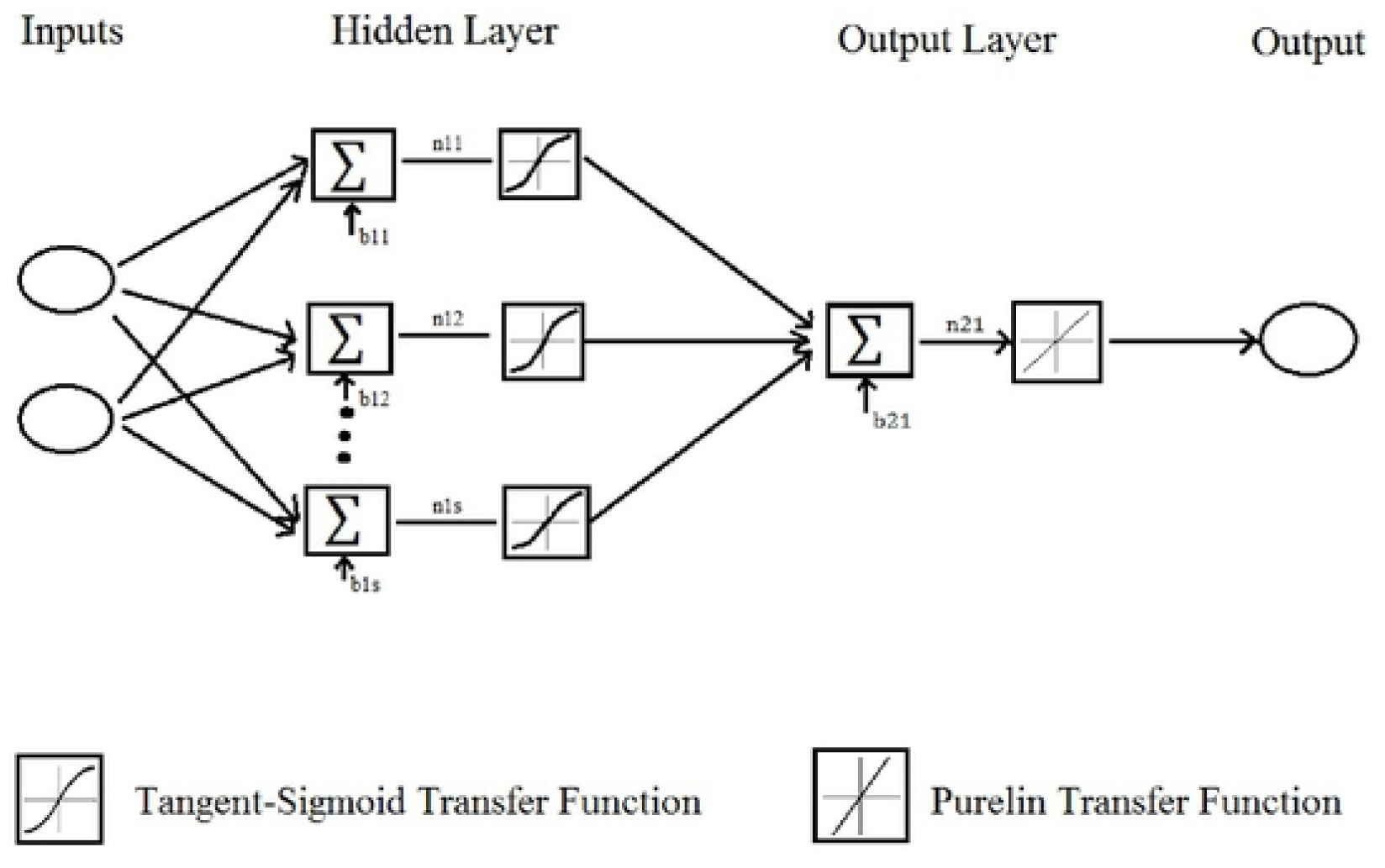
Structure of MLP ANN for modeling the microbial count. n, neuron; s, number of neurons; b, bias.

For each microorganism, a separate optimal ANN was developed. The training algorithm of ANNs was Levenberg-Marquardt (LV) backpropagation (BP). The transfer function in the hidden layer was Tangent-Sigmoid (tansig) and for the output layer, it was linear. Three replicates were carried out for each experimental run. Two statistical error criteria were taken into account for checking the goodness of fit for each ANN topology. These criteria were the coefficient of determination (R^2^) and root mean square error (RMSE):

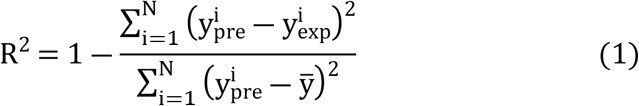

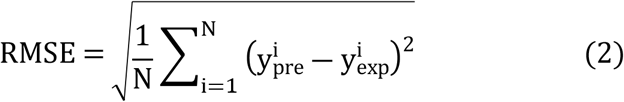

where, y_pre_ and y_exp_ indicate the predicted and experimental dependent variable (microbial count), respectively. ȳ indicates the average of the experimental dependent variable and N is the total number of runs. Based on the maximum R^2^ and minimum RMSE, the best ANN topology was selected. MATLAB (version 2013a, MathWorks, Massachusetts, United States) software was employed in ANN modeling.

### Statistical analysis

On each sampling occasion, a minimum of three observations was collected unless specified otherwise. Descriptive statistics of analysis results were calculated for each treatment. All data were initially evaluated by analysis of variance (ANOVA). The data were tested for homogeneity of variances at a significant level of P <0.05 and a probability value less than 0.05 was considered as statistically significant (Duncan, 1955). Microsoft Windows Excel 2010 and SPSS software (Version18.0, SPSS Inc., Chicago, USA) were used to analyze the data.

## Results and discussion

The effect of the temperature on the survival of microorganisms is seen in Fig. 2. For total viable count (TVC), the critical temperature in which the bacterial load was undetectable was 75 °C. There was not a significant difference (P>0.05) between control and samples processed at 60 °C in terms of the bacterial load. For lactic acid bacteria (LAB) and Enterobacteriaceae, the process temperature of 65 °C was considered as the critical temperature while this value for Psychrotrophs and Pseudomonas was 70 °C. Similarly, the process temperature of 60 °C was not effective in the inactivation of LAB, Enterobacteriaceae, Psychrotrophs, and Pseudomonas. Fig. 3 shows the rate of microbial growth in treated samples during storage at 4 °C. Although the critical temperature for TVC was 75 °C, microorganisms started to grow in samples treated at this temperature and the final population reached about 4 log CFU g^−1^ after three weeks of storage. But, according to the recommended limit of TVC which is 7 Log CFU g^−1^ samples treated at this temperature are still in the range of acceptable limits. This is also true for samples treated at 70 °C. The microbial load could be influenced by the P. psychrophila, A. allosaccharophila, and S. putrefaciens, which may be the most abundant bacteria as reported [19] from spoiled silver carp samples. The rate of Enterobacteriaceae growth in samples treated at 65 °C (critical temperature for Enterobacteriaceae) was very high, probably due to the incomplete inactivation of these microorganisms at this temperature. Even at higher process temperatures, the population of Enterobacteriaceae reached about 4 log CFU g^−1^ after three weeks of storage at cold temperatures. However, the presence of Enterobacteriaceae was not detectable in samples treated at 70 °C after one week and this period was two weeks for samples treated at 75 °C. These results are not far from those reported in common carp (*Cyprinus carpio*) where Enterobacteriaceae were detected after 14 days of storage in treated samples with sauce (30% tomato paste, 20% lemon juice, 30% oil, 10% garlic, 4% water, 3% salt, 1% red pepper, 1% cumin, and 1% thyme) and steam oven (90 °C, 15 min). LAB did not grow in samples treated at 75 °C even after three weeks of storage [20]. But for samples processed at the critical temperature of these microorganisms, the final population reached about one log CFU g^−1^ at the end of the storage period. These results are contradictory to some studies that detected LAB bacteria in considerable amounts in high treated samples (90 °C) using sous-vide processing technology in common carp (*Cyprinus carpio*) [19]. Similarly, a considerable growth of LAB bacteria in vacuum-packaged silver carp (*Hypophthalmichthys molitrix*) fillets were showed at 80 °C & 98 °C [21]. Psychrotrophs and Pseudomonas were not detectable in samples treated at 75 °C after two weeks of storage, however, their final population reached 3 and 2 log CFU g^−1^ at the end of the storage period, respectively. The growth of TVC, psychrotrophs, pseudomonas, and Enterobacteriaceae at high heat-treated samples (75 °C) could be associated with the thermolabile characteristics of these bacteria [22]. As reported, the growth of TVC in high treated samples (80 & 98 °C) was predominant in salted and vacuum-packaged silver carp (Hypophthalmichthys molitrix) fillets during storage [21]. Besides, TVC showed more heat resistance than Enterobacteriaceae in sous-vide cooked salmon loins processed by high pressure [4]. The overall results of microbial quality indicate that sous vide processing of silver carp at 75 °C was adequate for assuring the microbial quality of samples for two weeks. As it is seen in Fig. 3, the presence of microorganisms was not detectable at severe processing temperatures for a specific period (varied depending on the process temperature and type of microorganism) but, after that, they started to grow. This behavior was due to the recovery of thermally injured cells. The growing behavior of injured cells was different than viable cells in three ways: Firstly, the lag period of injured cells was increased considerably even using the optimum recovery medium. Secondly, the generation time of the injured cell can be higher than the total viable cells, even in the same intrinsic and extrinsic situations [23]. Thirdly, the recovery of injured cells was affected by the incubation temperature.

**Fig. 2.**
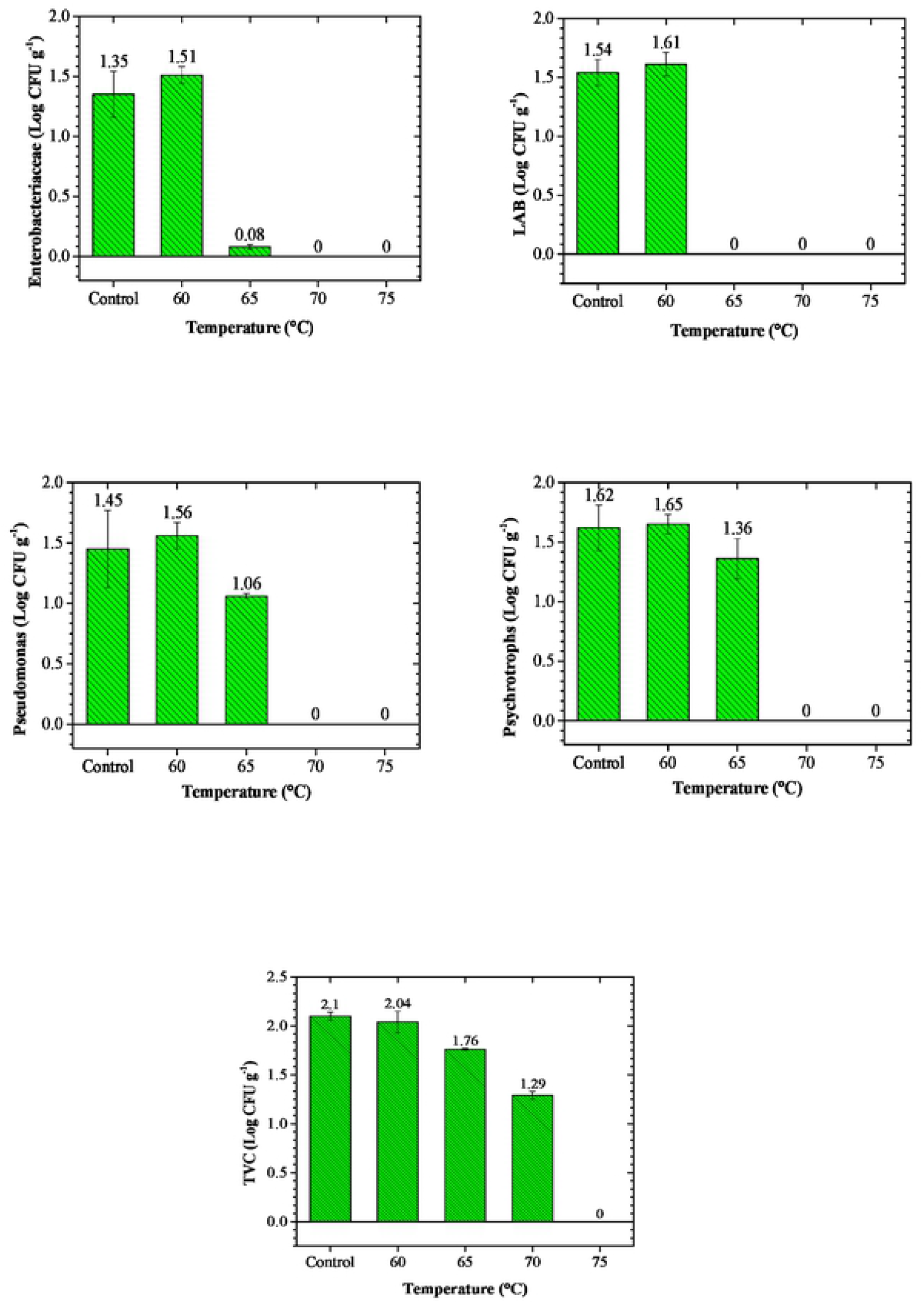
Results of microbial count (Enterobacteriaceae, Lactic Acid bacteria (LAB), Pseudomonas, Psychrotrophs, and total viable count (TVC)) of control and processed samples.

**Fig. 3.**
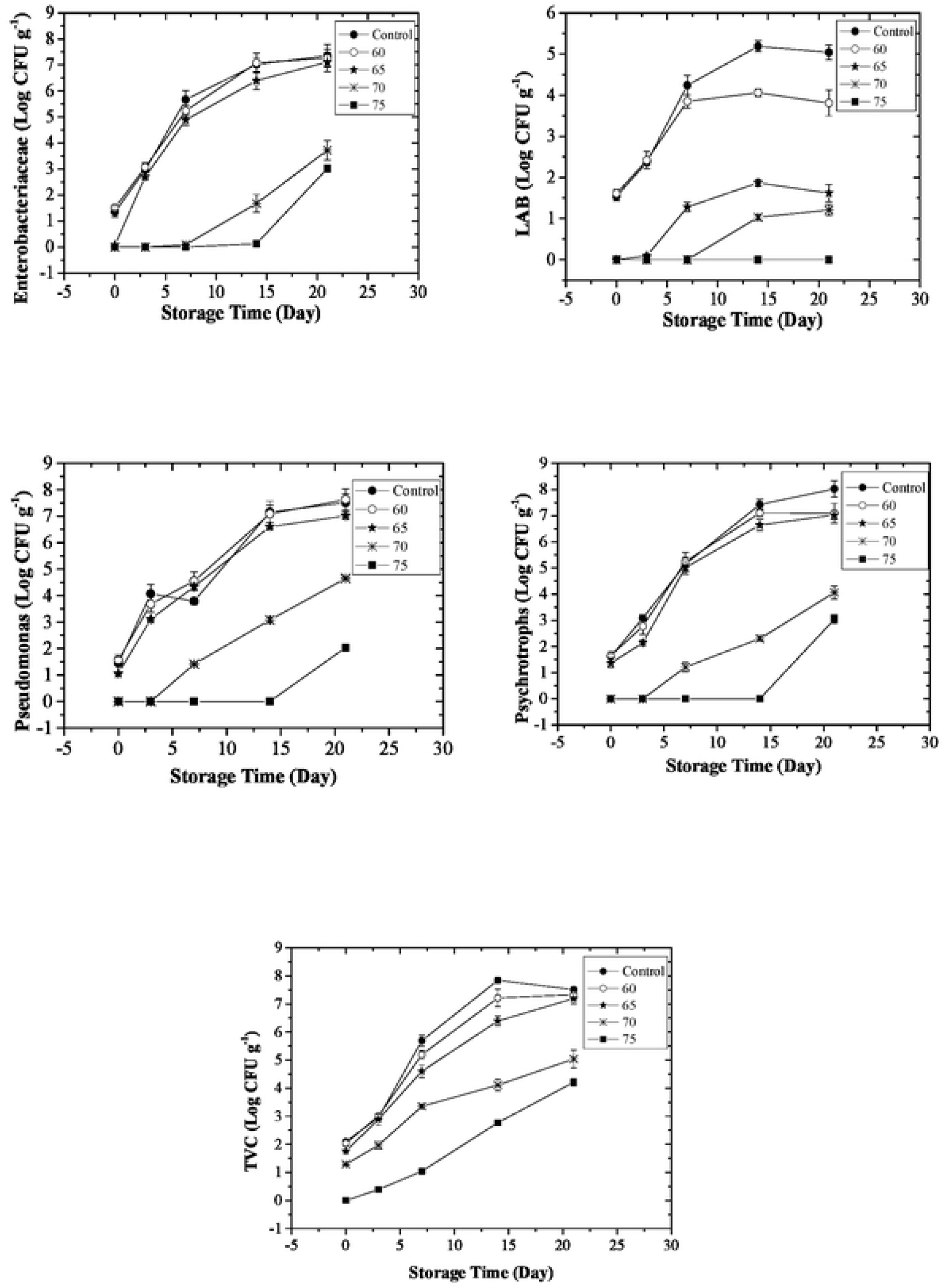
Growth kinetics of Enterobacteriaceae, Lactic Acid bacteria (LAB), Pseudomonas, Psychrotrophs, and total viable count (TVC) in control and processed samples during cold storage.

In the next step, an artificial neural network (ANN) was employed to build the model of the microbial quality which was affected by the process temperature and storage time. Process temperature (°C) and storage time (day) were considered as independent variables and the microbial load of the bacteria was chosen as the dependent variable. For each group of bacteria, a separate ANN was developed. For developing an optimum ANN for the bacteria, a different number of neurons (starting from 1 to 15) in the hidden layer were determined. Therefore, 15 networks, each containing a variable number of neurons in the hidden layer (1 to 15) were analyzed. The accuracy of each network was monitored based on the R^2^ and RMSE. One of the important issues that should be considered in ANN modeling is that if it is obtained a specific result in the first run, it will not necessarily get the same result in the next replicate [10]. This issue is due to the random selection and allocation of the input data for each section of the network development process, namely training, cross-validation, and testing. Therefore, it is recommended to repeat the run and report the average R^2^ and RMSE for each network with a different number of neurons in the hidden layer. In this study, the run was repeated five times, and, based on the average R^2^ and RMSE, the best ANN was obtained for each group of bacteria (Fig. 4).

**Fig. 4.**
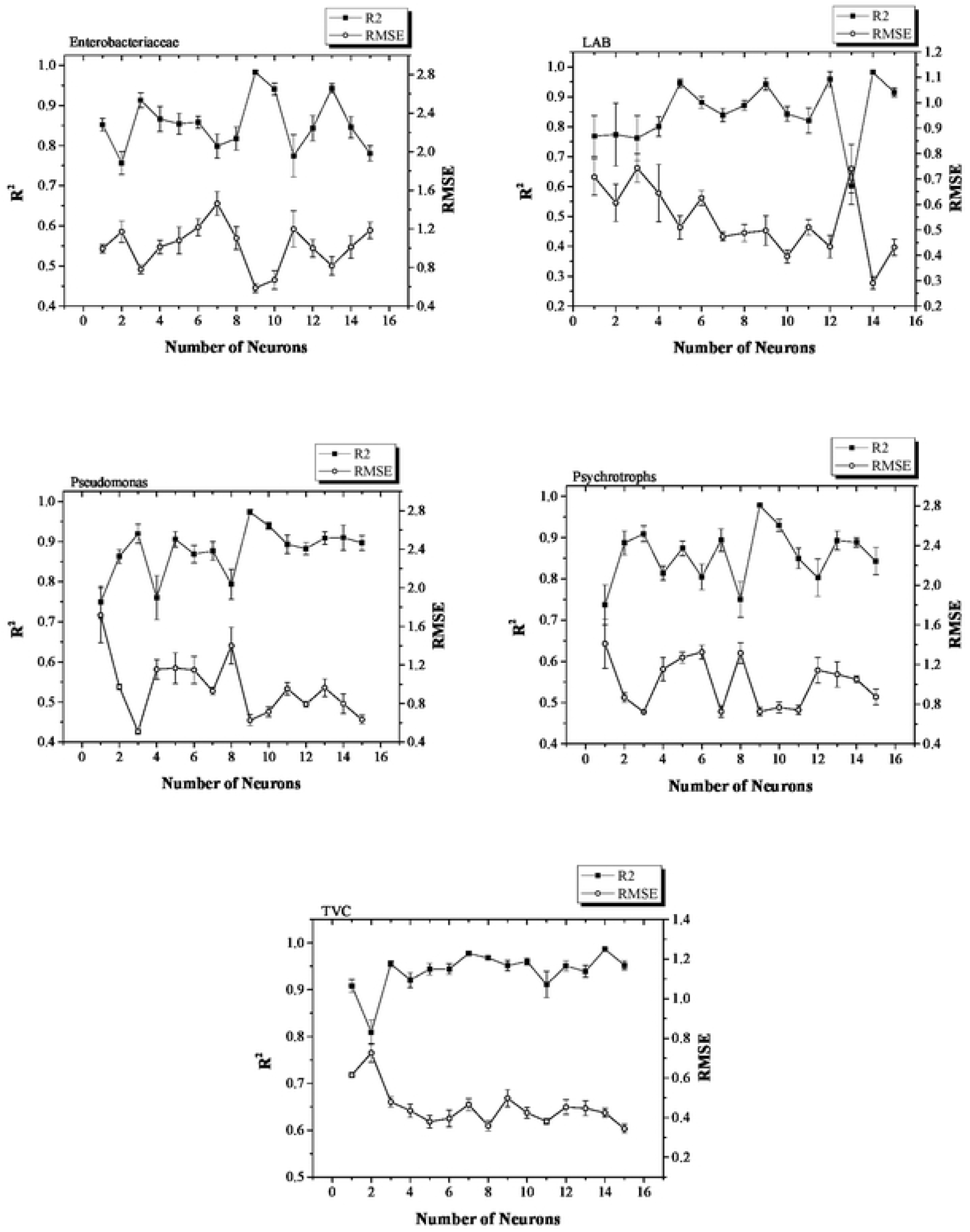
Average and standard error results of R^2^ and RMSE for five software runs in the procedure of finding the best number of neurons in the hidden layer of feedforward backpropagation ANNs.

For Enterobacteriaceae, Pseudomonas, and Psychrotroph the best ANN was obtained when the number of neurons in the hidden layer was 9, but, for LAB and TVC, this was 14 neurons. Fig. 5 shows the correlation between predicted (by the optimum network) and experimental data. As it is seen, a high correlation was obtained for all sets of bacteria which indicates the capability of ANN in modeling and predicting the microbial quality of silver carp processed by the sous-vide technique. This is especially important in industrial applications where the determination of optimum process conditions and prediction ability results in reduced costs and therefore higher profit.

**Fig. 5.**
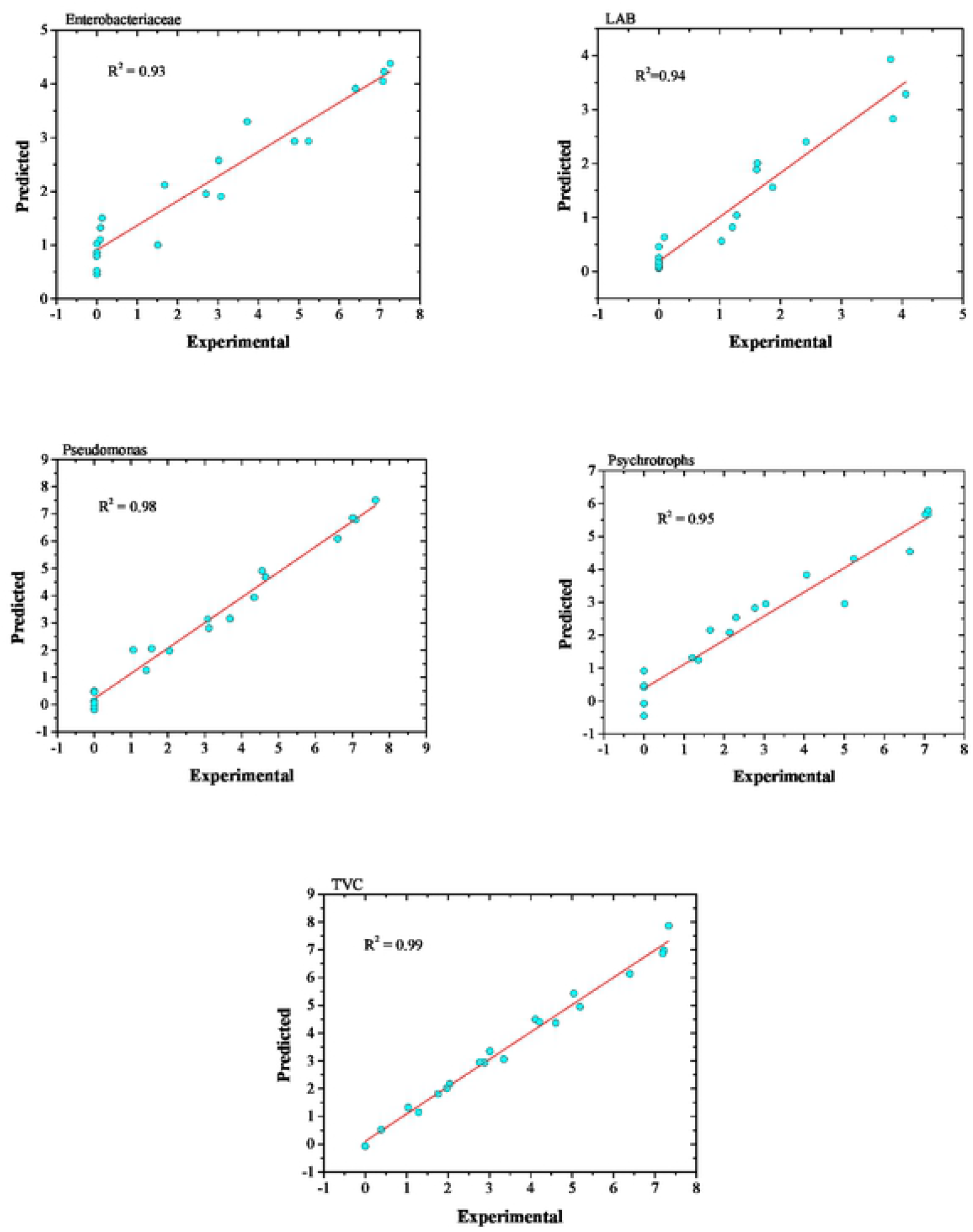
Comparison between predicted and experimental microbial count (Enterobacteriaceae, Lactic Acid bacteria (LAB), Pseudomonas, Psychrotrophs, and total viable count (TVC)). Predicted microbial counts were obtained by the best ANN topology.

Simulated results of ANN were used to generate the contour plots of microbial growth during storage (Fig. 6). These graphs provide useful and comprehensive information, especially for optimization purposes. From these graphs, for each group of microorganisms, the microbial load in silver carp fillets processed at a temperature between 60 to 75 °C storage up to 21 days was obtained.

**Fig. 6.**
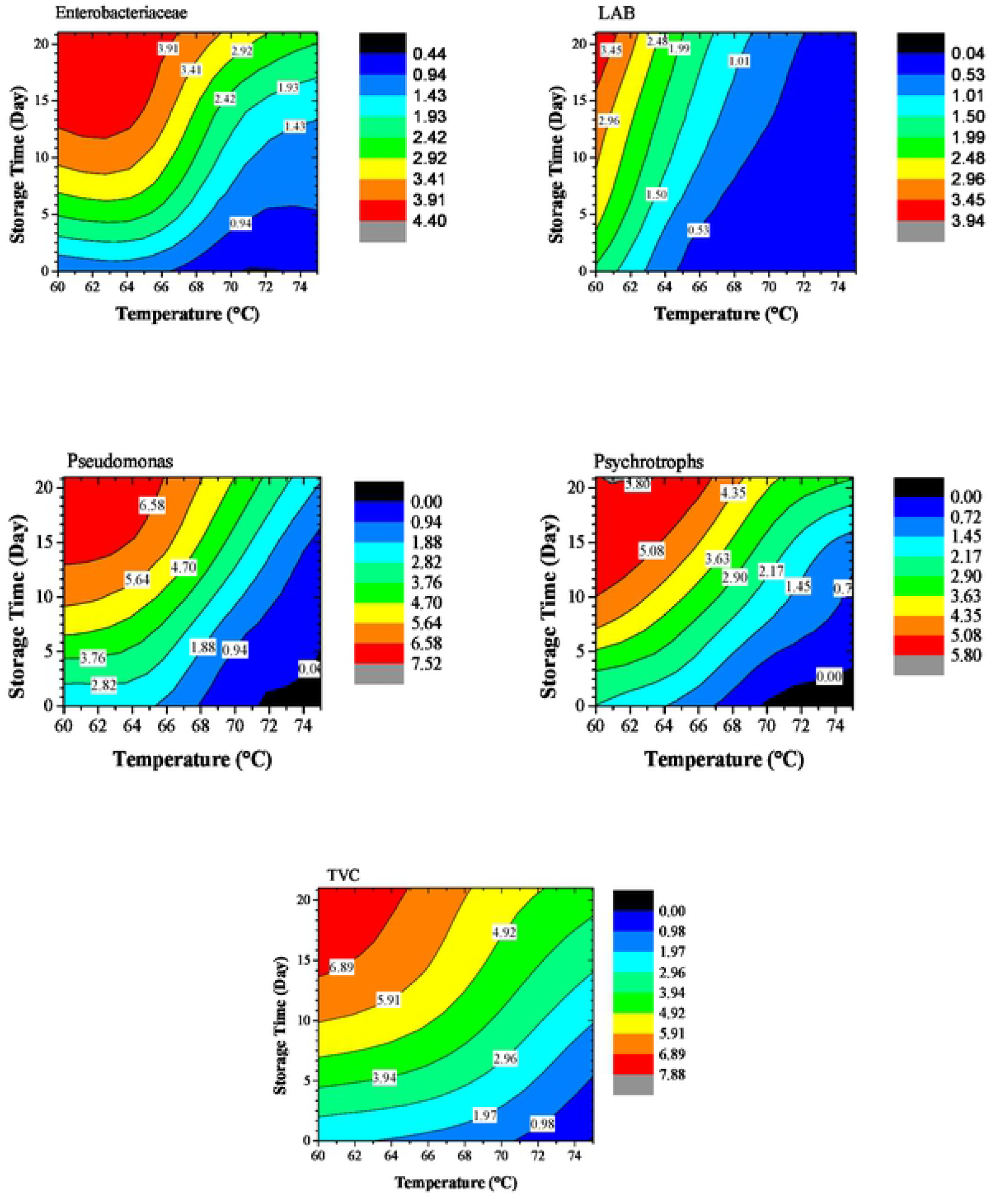
Simulated microbial count (Enterobacteriaceae, Lactic Acid bacteria (LAB), Pseudomonas, Psychrotrophs, and total viable count (TVC)) of sous vide processed samples at different temperatures during cold storage.

## Conclusion

This study reported the influence of sous vide processing on the effects of storage and processing temperatures on the silver carp (*Hypophthalmichthys molitrixi*) by analyzing the microbial load of LAB, psychrotrophs, pseudomonas, and Enterobacteriaceae. The results illustrated that sous vide processing at a higher temperature could significantly inhibit the microbial load of silver carp fillets. Moreover, the effects of process conditions and storage period on the microbial quality of samples were successfully modeled by artificial neural networks. This provides a powerful tool for manufacturers and researchers for the optimization of processes in which there is a complex relationship among variables. The simulation results of this research, presented by contour plots, provide a useful tool in optimizing the sous vide processing of silver carp, and also they can be used in designing an experiment for their studies of silver carp processing.

